# A proposed framework for evaluating meat alternatives

**DOI:** 10.1101/2024.11.18.624184

**Authors:** Cameron Semper, Caroline Kolta, MJ Kinney, Jordan Giali, Michaella Rogers, Dan Blaustein-Rejto, Amy C. Rowat, Olivia J. Ogilvie, Ryan Hutmacher, Josephine Wee, Isaac Emery, Laura J. Domigan, Kantha Shelke

**Affiliations:** Department of Microbiology, Immunology and Infectious Diseases, University of Calgary, 3330 Hospital Drive, Calgary, Alberta, T2N 4N1, Canada; XPRIZE Foundation, 10736 Jefferson Blvd, Culver City, California, 90230, United States; The Breakthrough Institute, 2054 University Ave, Berkeley California, 94704, United States; Department of Integrative Biology & Physiology and the Rothman Family Institute for Food Studies, University of California, Los Angeles, Los Angeles, California, 90095, USA; School of Biological Sciences, University of Canterbury, Christchurch 8041, New Zealand; Biomolecular Interaction Centre, University of Canterbury, Christchurch 8041, New Zealand; Well Beyond Food, LLC, South Portland, Maine, 04106, USA; Department of Food Science, The Pennsylvania State University, University Park, PA 16802, USA; One Health Microbiome Center, Huck Institutes of the Life Sciences, University Park, PA 16802, USA; WSP USA Inc., 1001 4th Ave, Suite 3100, Seattle, WA 98154; Department of Chemical and Materials Engineer, University of Auckland, Auckland, New Zealand; Regulatory Science and Food Safety Regulation, Johns Hopkins University, Washington DC, 20001, United States

## Abstract

Concerns surrounding the environmental, economic, and ethical consequences of meat production and industrial agriculture have prompted substantial research and capital investment into the production of meat alternatives. Alternative meat production encompasses a variety of technological approaches including plant-based meats, cell-based or cultivated meats, meat alternatives relying on fungal protein sources, and hybrids thereof; each of which offers unique advantages and disadvantages and has been associated with a myriad of claims supporting it as the preferred alternative to animal-derived meats. As part of XPRIZE Foundation’s Feed the Next Billion competition, we developed a framework for evaluating meat alternatives by measuring their structural, nutritional, and organoleptic properties while also assessing safety and their purported environmental and economic benefits compared to animal-derived meats. The framework is technologically agnostic and can be used to evaluate meat alternatives of all types. The output of the framework enables a data-driven comparison to animal-derived meat and/or other alternative meats, allowing a range of stakeholders (e.g., food startups, investors, government) to assess technological readiness, competitive advantage, and impact potential. This framework can assist this nascent industry as it moves towards standardizing approaches to evaluating the quality, safety and proposed benefits of meat alternatives.

## Introduction

Climate changes, population growth, and increasing demand for meat coinciding with economic development ^1^ have placed existing agricultural systems under incredible strain. These pressures have renewed efforts to examine the health and sustainability challenges associated with traditional livestock farming ^2^. Examples of the complex interplay between agricultural systems and human health include the proliferation of antimicrobial resistance stemming from the overuse of antibiotics and the emergence of zoonotic diseases ^3^, such as avian influenza ^4^. Agriculture is also a significant contributor to global anthropogenic greenhouse gas (GHG) emissions ^5,6^, with a recent study finding that rapid phase-out of animal agriculture has the potential to stabilize GHG levels ^7^ and help achieve the goals of The Paris Agreement ^8^. These are challenges, along with deforestation ^9^, fresh water use ^10^, and biodiversity loss ^11^, that risk being exacerbated by increased livestock production. They are also motivations behind the burgeoning field of ‘alternative proteins’ or ‘cellular agriculture’, which seeks to develop complementary approaches to producing animal meat or protein rich alternatives that can serve as faithful replacements for animal origin (AO) products. Environmental modeling has predicted or shown that these alternatives can serve as a more sustainable source of dietary protein compared to traditional animal agriculture ^12,13^, while providing other benefits such as a reduction in antibiotic usage and improvements to animal welfare ^14^.

The history meat alternatives dates as far back as the 6^th^ century ^15^ and includes a variety of plant-based examples such as tofu, tempeh and seitan. Life cycle assessments have found that many plant-based meat alternatives have a lower environmental impact than animal-derived sources of dietary protein ^13^; however, these products generally do not closely replicate the organoleptic qualities of conventional meat. Academic and industry led efforts have focused on developing meat alternatives that more closely replicate these qualities, as exemplified by Dr. Mark Post’s cell-based hamburger ^16^ and the commercialization of Beyond Meat and Impossible Food’s plant-based burgers, sausages, and nuggets. These examples are representative of the majority of those currently available to consumers – alternative products that aim to replicate ground or minced meats. More recently, whole-cut alternative meat products have come to market including plant-based salmon in parts of the EU and hybrid chicken alternatives composed of plant-based and cell cultured ingredients in Singapore ^17^. Release of these products has coincided with economic modelling highlighting outstanding technical challenges surrounding certain approaches to alternative meat production ^18,19^, and consumer adoption remains heavily impacted by pricing, organoleptic qualities, and product familiarity ^20^. Combined with a number of companies in the industry missing projected release dates for their own meat alternatives ^21^ and declining sales of existing products on the market ^22^, it is clear that alternative meat production requires substantial innovation, investment, and maturation before the proposed benefits to environmental sustainability and human health can be achieved.

In effort to promote innovation and accelerate technological maturation of alternative meat production, the XPRIZE Foundation launched Feed the Next Billion (FTNB), a USD $15 million prize competition focused on incentivizing competing parties to develop structured chicken breast or fish fillet alternatives. The alternative meats produced were tasked with replicating or outperforming conventional chicken or fish across a variety of metrics including access, environmental sustainability, animal welfare, nutrition, taste, and texture. The competition was solution/technology agnostic and invited submissions incorporating a variety of alternative protein sources including plant-based, cell-based or cultivated (henceforth referred to as ‘cell-based’), mycoprotein and single-celled protein (SCP), or hybrids of these alternatives. The multi-year competition curated a set of established methodologies to assess the characteristics of the competition submissions which form the foundation of a framework that was developed for evaluating meat alternatives. This framework was designed to comprehensively assess meat alternatives using methodology that was robust, reproducible, and cost effective. The results obtained through application of this testing framework were used as the basis for determining advancement throughout the various stages of the competition. We successfully applied our framework to plant-based, cell-based, and mycoprotein-based chicken and fish alternatives. Importantly, the framework allowed for meaningful assessment of meat alternatives compared to the animal-based product they aimed to replicate, as well as to other alternative products in the competition. Furthermore, assessment was possible across different product types (i.e., alternative chicken breast vs. alternative fish fillet) and product technology (i.e., plant-based vs. cultivated), resulting in a generalized approach to evaluating meat alternatives.

### XPRIZE Feed the Next Billion Framework

The framework was designed to assess both product and process level parameters of meat alternatives (Figure 1). The portfolio of tests included within product level assessment employs methodologies with an established history of use in food analysis. Collectively, these tests measured the size, structural and physical characteristics, and nutritional profile of the meat alternatives within the competition. Additional product level assessment looked at the sensory characteristics and organoleptic properties of meat alternatives using consumer sensory panels, as well as an assessment about product versatility with respect to preparation and cookability. Throughout the competition, all AOR products were subjected to the same testing framework, resulting in generation of a baseline data set used to make comparisons between meat alternatives and the animal-derived products they sought to imitate.

**Figure 1.**
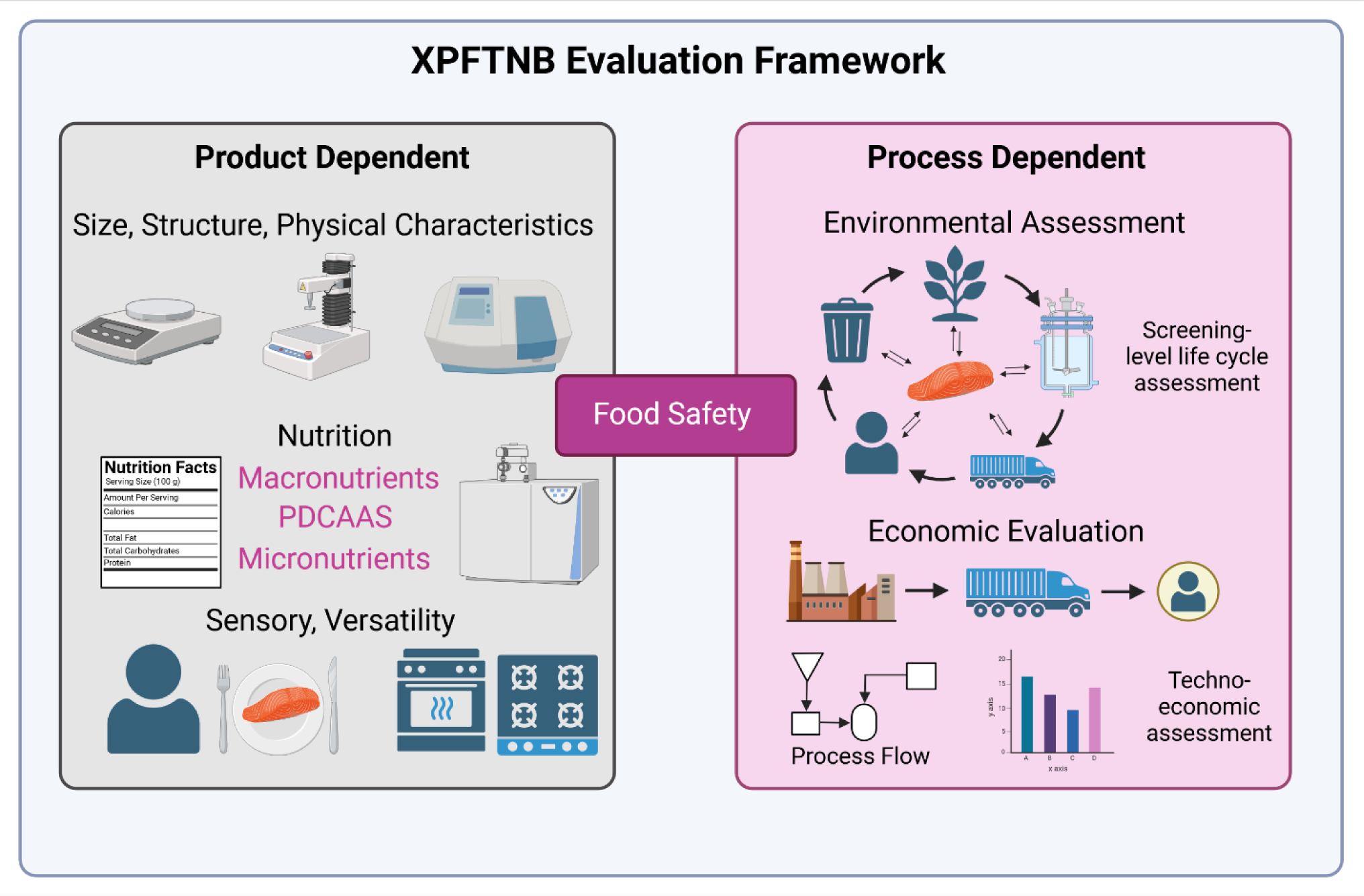
Framework Overview.

Process level assessment focused on two main areas, the environmental sustainability and economic feasibility of the meat alternatives. Broadly speaking, this assessment included a screening-level life cycle assessment (LCA), and a techno-economic assessment (TEA) based on data specific to the processes used to produce the meat alternatives. These assessments incorporated information related to ingredients and processing aids, manufacturing and storage, as well as distribution, market launch strategies and regulatory considerations. For the environmental and economic assessment, existing data related to AOR products was gathered from available databases and used for comparison where appropriate.

A key component of this framework that considered product and process level factors was food safety. Process level factors were considered upstream elements and were reviewed throughout the duration of the competition. This review focused on risk identification, mitigation and management to ensure samples were safe for consumer testing without requiring a formal regulatory filing. Product level factors encompass more traditional food safety testing to ensure the absence of microbial and chemical contamination that could acutely compromise food safety.

### Product Level Assessment

#### Structure and Physical Characteristics

The texture, mouthfeel and overall organoleptic properties (consumer experience) of animal meat are influenced by the structure and physical characteristics of the tissue or food matrix, which in turn, is dependent on factors such as the species, tissue type, post-mortem handling ^23^, and food processing. Producing structured meat alternatives is inherently more challenging than ground or unstructured alternatives ^24^. In the case of plant- and fungal-derived proteins, the manufacturing process typically involves techniques, such as extrusion, to modify texture and mimic the fibrous nature of muscle ^25^ . For cell-based alternatives, mimicking the microstructure of meat can involve controlling cell differentiation (e.g., myogenesis) and other approaches conscripted from the field of tissue engineering ^26^. Faithfully replicating the properties of whole-cut meat is seen as critical for the success of alternative meats; whole-cut meat products account for the largest proportion of retail sales by dollar volume, and are preferred by consumers over ground products ^27^. For these reasons, XPRIZE Feed the Next Billion required submissions to replicate whole-cut chicken or fish, and relied on a collection of robust, well characterized approaches for characterizing the macrostructure of meat to assess performance.

#### Size

Assessing the mass of submissions ensured they conformed with the standard serving size (85-115g) of conventional meat. This requirement was implemented to increase the challenge associated with developing whole-cut or fillet alternatives while promoting development of products that aligned with consumer expectations. Mass measurements were undertaken using a laboratory scale, with a minimum of 15 samples measured (out of a total of 200+ samples).

Submissions were not required to have a uniform mass, but all samples needed to be a minimum of 85 g within the competition to be considered compliant. Submissions were also required to have a minimum thickness of 1 cm. The implementation of this parameter was to disincentivize the development of thin, elongated submissions that met the mass requirements but would otherwise be considered “atypical” compared to AORs. Structured cell-based meat samples that have thickness akin to conventional cuts of meat are still in development ^28^, though proof-of-concept studies show promise for future development of thicker, structured cell-based meats ^29^. The minimal thickness requirement was also included to ensure submissions were compatible with other testing methodologies utilized within this framework (discussed below).

#### Colorimetric testing

Visual appearance is a major determinant of purchase intent for meats with color being one of the major parameters used to gauge quality ^30^, safety, freshness ^31^, and consistency ^32^. This is exemplified by certain industry practices, such as the addition of carotenoids to the diets of farmed salmon to imbue them with the pink-red coloration typical of wild salmon ^33,34^. Color is also an important indicator of the cooking process where it is used as a proxy for food safety and desired organoleptic properties ^35^. Within this framework, color was assessed before and after cooking and compared to AOR samples in the equivalent state. Colorimetric similarity to AORs prior to cooking did not strongly correlate with similarity post cooking. This could be due to the heat labile nature of certain components used in alternative meat development and highlights a need for continued development in the field.

#### Water holding capacity (WHC)

Water holding capacity (WHC) refers to the ability of meats and meat alternatives to retain their internal fluid ^36^. Several approaches to determining WHC exist, and a common approach applies low-speed centrifugal force to samples and measures the amount of fluid released. WHC has traditionally been used as an indicator of meat quality, with the internal water content contributing to perceived “juiciness” of meats during chewing ^36^. WHC has also been shown to correlate with visual acceptability of meat products, highlighting its importance as a tool for assessing overall quality ^30^.

WHC is a key characteristic of meat quality, and it has been applied as a tool for evaluating meat alternatives in recent years ^30^. Though the correlation between WHC and cook loss is imperfect, it can nonetheless be an indicator of the ability of foods to retain moisture during the cooking process. For XPRIZE Feed the Next Billion, WHC was assessed on each submission AOR prior to cooking and post cooking. Structural changes that occur during the cooking process, due in part to heat-induced protein denaturation, are known to impact the WHC of animal-based meats ^37^. In contrast to the observations during colorimetric testing, WHC testing showed a strong correlation between results obtained prior to cooking and post cooking. Submissions that produced WHC values similar to their respective AORs in pre-cooked state were more likely to have similar WHC to the AOR in a post-cooked state (Table 1).

**Table 1.** Pre and post cook correlation of WHC testing results of meat alternatives (results have been anonymized)

| **Competition Ranking** | **WHC, relative to AOR (pre-cook)** | **WHC, relative to AOR (post cook)** |
| --- | --- | --- |
| **1** | Submission 1 | Submission 1 |
| **2** | Submission 2 | Submission 2 |
| **3** | Submission 3 | Submission 4 |
| **4** | Submission 4 | Submission 3 |
| **5** | Submission 5 | Submission 6 |
| **6** | Submission 6 | Submission 5 |

#### Texture Profile Analysis (TPA)

TPA, also known as the double compression test, utilizes a texture analyzer to compress standard sized cores cut from a given sample. The output of this test is a force/time curve, from which several characteristics are derived including cohesiveness, springiness, chewiness, hardness, and resilience ^38^. TPA is intended to test samples under conditions that replicate the chewing process, and compared to Warner-Bratzler Shear Force testing, has correlated better with sensory analysis ^39^. TPA has been used widely in the field of meat science, including for texture analysis of whole-cut ^40^ and ground meat products ^41^. It can be used to analyze the texture of uncooked and cooked samples and has been successfully used to assess meat alternatives ^42^, including cell-based meat ^43^. Meat texture contributes significantly to the overall sensory experience, indicating it will be an important parameter to optimize for meat alternatives. TPA allowed for comparison of submissions to their AOR across specific parameters that influence overall texture.

### Warner-Bratzler Shear Force (WBSF)

WBSF has traditionally been used to assess meat tenderness, which is considered an indicator of product quality and a contributor to the sensory experience. WBSF testing has been applied to whole muscle meat products ^40^, sausages and other ground meat products ^44^, as well as meat alternatives ^45^, highlighting its utility in assessing textural parameters. WBSF has been described as measuring the hardness of meat ^46^, and in our testing, we saw strong correlation between WBSF results and the hardness results obtained through TPA.

### Nutrition

XPRIZE Feed the Next Billion prioritized nutritional equivalence as a key performance indicator throughout the competition. Nutritional assessment generally followed the US Food and Drug Administration (FDA) Nutrition Facts label and considered key macronutrients (e.g., protein, fat) as well as an analysis of micronutrients. The competition framework promoted development of submissions that replicated or improved upon the nutritional profile of conventional chicken or fish. Improvement could be through increasing the quantity of desirable nutrient (e.g., protein, polyunsaturated fats) or eliminating anti-nutritional factors such as mercury found in marine and some freshwater fish ^47^.

### Protein content and quality

Meat is a major source of dietary protein, and its protein content is a significant factor influencing consumer choice ^48^. The XPRIZE Feed the Next Billion framework measured protein content and protein quality during its assessment. Standardized AOAC methodology ^49^ was used to determine the protein content per serving size, which was compared to the respective AOR for each submission. The FTNB framework utilized *in vitro* Protein Digestibility Corrected Amino Acid Score (PDCAAS) testing to determine protein quality of each of the competition submissions to facilitate comparison to AORs ^50^. While PDCAAS does not account for anti-nutritional factors ^51^, it remains a well-established approach to gauging protein quality. Assessment of protein quality is especially important in the context of alternative proteins. Many sources of plant-based proteins contain limited amounts of essential amino acids ^52^ necessitating the use of amino acid supplements or blending two or more plant protein sources with complementary amino acid profiles ^53^ to overcome these limitations and achieve a complete protein source.

### Fats

Fats are a critical macronutrient found in meat and an important contributor to the flavour, texture and overall organoleptic profile. The fat content in meat varies widely, and some sources of animal-based meat are sought after for their high content of desired fats. Fish consumption is recommended in part due to its high content of polyunsaturated fatty acids (PUFAs) ^54^. When developing meat alternatives, the specific fat profile needs to be taken into account to ensure nutritional equivalence while meeting consumer expectations. The XPRIZE Feed the Next Billion framework assessed total fat (g/serving size) for each of the submissions, while also requiring that submissions aiming to replicate fish AOR contain comparable levels of PUFAs.

### Carbohydrates

Meat typically contains very low levels of carbohydrates making true nutritional equivalence difficult to achieve for some types of meat alternatives. Plant-based protein sources often contain carbohydrates, and sugars have been used in meat alternatives to imbue them with the ability to brown during cooking or for flavor enhancement. For the FTNB competition, submissions were subjected to sugar profiling by ion chromatography ^49^. Sugar levels above 0.5 g per serving size were considered as having the potential for “added sugars”. Dietary fibre was also assessed within the FTNB framework; however, deviation from the AOR with respect to fibre was not considered when determining overall nutritional equivalence. The inclusion of fibre has been shown to provide structural advantages when using extrusion technology ^55^ and dietary guidelines encourage fibre intake ^56^. Indeed, it has been argued the presence of fibre in meat alternatives improves their overall nutritional profile compared to conventional meat ^57^.

### Sodium

The high sodium content of plant-based foods has been a major criticism to-date ^58–60^, and for meat alternatives to offer nutrition that replicates or outperforms conventional meat, the sodium content will need to align with dietary recommendations. Accordingly, submissions were encouraged to match the sodium content of their AOR. Further guidance was provided for submissions to keep sodium content below 300 mg/serving size and thereby under the threshold (20% of daily value) where foods become categorized as ‘high in sodium’ by many dietary guidelines.

### Micronutrients and cholesterol

Micronutrient assessment focused on minerals included on the US FDA Nutrition Facts label as well as vitamin D and vitamin B12. Iron, potassium and calcium are important dietary components, with deficiencies contributing to adverse health effects such as anemia ^61^ and loss of bone density ^62^. Vitamins B12 and D are essential vitamins and products of animal-origin are a major dietary source of them ^63^. The FTNB framework included these micronutrients to foster the development of meat alternatives that were nutritionally complete and able to fully replace their respective AORs from a nutritional standpoint.

Cholesterol was assessed for submissions incorporating animal cells in their product formulation. Submissions were required to keep cholesterol levels below 110% of the AOR to be considered nutritionally equivalent. The baseline amounts of cholesterol found in cell-based foods have not been well established, and guidelines surrounding cholesterol content will require adjustment as more data becomes available.

### Product Versatility

Meat is well regarded for its ability to withstand a myriad of preparation techniques. For a meat alternative to replicate or outperform an AOR, it must display comparable levels of cooking versatility and be capable of maintaining a cohesive structure and appearance across a range of culinary applications. The FTNB framework assessed versatility by subjecting submissions to multiple cooking techniques by culinary professionals. The cooking preparation methodologies assessed included dry heat cooking, moist heat cooking, dry heat cooking in combination with breading, and denaturation methodology incorporating the use of acid marination. The texture and performance of the submissions under each of the cooking techniques was rated as well as the overall ease of use of the product from the standpoint of a chef or individual using the meat alternatives in meal preparation. The rationale behind selection of the preparation conditions tested was based on common uses of chicken and fish. These meats are commonly breaded and cooked using dry and moist heat methods. Acid marination is another popular preparation technique applied to chicken and fish; however, it was also included to assess the stability of competition submissions under low pH conditions. Meat alternatives commonly incorporate binding and/or gelling agents such as methylcellulose and carrageenan that can be critical for their structure and texture ^64^. The stability and functionality of these agents can be compromised at low pH due to acid hydrolysis ^65^, potentially impacting the versatility of meat alternatives under acidic conditions.

### Sensory Profile

Many of the *in vitro* tests used evaluated specific aspects of meat alternatives that are important for the sensory experience; however, a consumer tasting panel is the ultimate measure of sensory performance and by extension, consumer acceptance. Achieving a sensory profile desirable to consumers and comparable to the AOR will be critical for widespread adoption of meat alternatives ^66^. Sensory analysis was therefore a fundamental aspect of the FTNB framework as it allowed for an assessment of submission performance under intended-use conditions. Sensory analysis also provides feedback from potential adopters of meat alternatives, allowing for product iteration and improvement based on key findings.

Within the FTNB framework, sensory profile analysis was performed using a 100-person consumer panel comprised of a proportional distribution of gender, age, income and nationality. Prior to undertaking the consumer panel, all submissions were subjected to food safety testing in accordance with local regulations (see ‘Food Safety’ for full description of safety testing and considerations within the FTNB framework). All participants selected for the consumer panel self-identified as non-rejectors of the AOR (i.e., consume chicken or fish with some regularity) and communicated a willingness to consume meat alternatives. The taste testing was blinded, and the competition submissions were evaluated ‘as-is’, as well as incorporated into recipes. Sensory testing followed a sequential monadic approach, and tasters were asked to rate competition submissions and AORs for overall liking, visual appearance and aroma, texture and mouthfeel, and taste and aftertaste. Tasters rated performance in each of these attributes using a scale of 1 to 7 (Very Poor - Excellent). In addition to the sensory attributes evaluated, tasters were also asked to make a holistic assessment of each of the submissions after the blinded taste test was completed. This involved grading the samples for how similar they were to their respective AORs using a five-point scale (Not close at all – Very close), as well as declaring purchase intent for the alternative meat products, ranging from Not Likely to Very Likely.

Comparison between the results of sensory analysis and of the structure and physical characteristics testing showed that TPA was the best predictor of performance in “texture and mouthfeel” compared to other tests analyzed (Table 2). We found that similarity to AOR in “Springiness” was the best correlator of performance in the texture attribute of sensory analysis, though additional analysis using larger data sets is required to statistically validate this observation and determine if it can be used as a reliable predictor of sensory performance for alternative meat products.

**Table 2.** Correlation between TPA-Springiness and texture assessment according to consumer sensory panel (results have been anonymized).

| **Competition**<br>**Ranking** | **Springiness,**<br>**relative to AOR**<br>**(cooked)** | **Overall texture,**<br>**sensory**<br>**evaluation** |
| --- | --- | --- |
| **1** | Submission 1 | Submission 1 |
| **2** | Submission 2 | Submission 2 |
| **3** | Submission 3 | Submission 3 |
| **4** | Submission 4 | Submission 4 |
| **5** | Submission 5 | Submission 6 |
| **6** | Submission 6 | Submission 5 |

### Process Level Assessment

#### Screening Life-Cycle Assessment (LCA)

Traditional and anticipatory LCAs of these products have indicated they have potential to substantially reduce land use, water use, and GHG emissions ^12^, while also helping to mitigate public health concerns related to zoonotic disease and antimicrobial resistance. The scale of the environmental benefit is influenced by several factors including the specific type of meat alternative (i.e., plant-based meat vs. cell-based meat) ^13^, specific production or formulation variables (i.e., plant protein source ^67^, growth media ^68^) as well as the species of animal meat (i.e., chicken vs. beef ^69^) and the production system (i.e., intensive vs. extensive ^70^) used for comparative studies. LCAs have consistently found net environmental benefits associated with substituting beef and other ruminant meats with alternative meats ^12,71–73^. When compared to chicken and fish, the results are more nuanced with alternative meat products offering environmental benefits in some categories but drawbacks in other impact areas ^73^. For example, alternative meats compare favourably to traditional chicken in land use but worse in energy use 73.

The FTNB framework incorporated screening-level LCAs to model the environmental footprint and overall sustainability of submissions. The results of the screening level LCA were used to compare the environmental footprint of meat alternatives across the measured impact categories to their respective AORs. Critically, the screening level LCAs were based on empirical data allowing for comparisons reflective of the highly specialized submissions and the processes used to produce them. Screening LCA data was also used to identify key product or process level ‘hot spots’ – major contributors to impact in specific categories. Hot spot identification allowed for product reformulation or process modification to help mitigate the environmental footprint of competition submissions ahead of final judging. Common process level hot spots identified included reliance on energy grids powered by non-renewables as well as transportation of ingredients. Product level hot spots included specific food ingredients and equipment required for cultivation of cells and/or microbes. The ability of competing teams to incorporate screening LCA results into their product and process design allowed for refinement with a focus on environmental sustainability. Product iterations that incorporated feedback from the screen LCA showed marked sustainability improvements.

### Techno-Economic Assessment (TEA)

The XPRIZE FTNB framework utilized techno-economic analysis to assess the commercial viability of submissions. This assessment looked at the supply chain, manufacturing and production processes, and organization structure of each of the competing teams. The input data was then used to model the technological and market readiness of each competitor. Process flow diagrams overviewing production were included and used to analyze system elements and boundaries. Competing teams were also asked to provide a go to market strategy including plans for scaling production of their meat alternatives. This information was used to forecast the profitability potential and the market impact of each of the submissions. Finally, the degree of uncertainty associated with the analysis was assessed and included in the overall economic evaluation of each competing team. TEA resulted in identification of ‘hot spots’ that could compromise or limit the economic viability of submissions. Economic hot spots included process and product level considerations, and teams were able to iterate and improve in response to hot spot feedback. The inclusion of economic modelling within the FTNB framework helped with assessing the potential of submissions to achieve the overall goal of the competition. Economic modelling also proved helpful from a product formulation standpoint, allowing teams to make decisions to maximize performance across all framework criteria.

### Food Safety

Alternative meat products are increasingly incorporating novel ingredients (e.g., recombinant proteins) and/or new manufacturing processes not previously applied in food production (e.g., cell culture for cell-based meat). These factors lead to many (but not all) alternative meat products being classified as novel foods. Consequently, the global regulatory environment for alternative meats remains nascent and food safety has been a salient topic for the field. Recent efforts led by the Food and Agriculture Organization of the United Nations (FAO) ^74^ have highlighted some unique food safety considerations associated with emerging technologies such as precision fermentation, single celled protein, and cell-based meat. XPRIZE FTNB featured submissions using a wide range of approaches to producing meat alternatives, creating a need for a bespoke approach to food safety to ensure submissions adhered to existing food safety regulations and best practices, while also accounting for the unique food safety concerns associated with novel foods. The FTNB framework approached food safety to account for process and product level considerations.

### Input-related safety concerns

All submissions to the FTNB competition were required to disclose a full list of ingredients detailing the supplier, country of origin, and declaring any known allergens. What qualifies as a known allergen is country-dependent ^75^, so teams were required to declare any ingredients listed as known allergens in the United States, Canada and their country of origin. Based on this list, follow up documentation or testing was required to account for the specific food safety concerns associated with different approaches to solution development. Food safety risks associated with plant-based proteins and other ingredients include heavy metal and mycotoxin contamination ^76^. To account for these, quality control (QC) documentation from plant-based protein/ingredient suppliers was required asserting the ingredients were free of mycotoxin and heavy metal contamination. In instances where supplier documentation was insufficient, could not be provided, or was outdated, the competition submissions were subjected to standardized laboratory testing for heavy metals and mycotoxins.

Assessment of ingredient and input lists involved categorizing them as having received approval from a national regulatory agency or lacking it. Regulatory approval was defined as ingredients having received Generally Recognized as Safe (GRAS) or Qualified Presumption of Safety (QPS) status or having a prolonged history of use in food applications. For these ingredients, links to a curated GRAS/QPS inventory and supplier QC documentation was included as part of the food safety review. For ingredients lacking regulatory approval, teams were required to perform a self-GRAS assessment and include supportive literature or documentation providing evidence of the safety of the specific ingredient. This information was reviewed by food safety, legal and food regulation experts prior to advancement in the competition.

The review of input-related safety concerns was an iterative process undertaken at multiple stages throughout the competition. This approach was beneficial to teams and to the competition, as any food safety concerns were identified proactively, allowing time for adjustment, product reformulation, or additional testing or disclosure. As the field of alternative meat continues to mature, we anticipate a commitment to food safety from the onset of product development will greatly enhance product quality and expedite navigation of regulatory requirements.

### Post-production product related safety concerns

Post-production food safety involved implementing best practices for the transport, storage and handling of competition submissions, as well as employing standardized food safety testing for detection of foodborne pathogens. Samples were transported frozen and competing teams were required to provide instructions for safe thawing and handling. All samples were cooked prior to serving, according to guidelines for their respective AORs. Chicken breast, and competition submissions aiming to replicate chicken breast, were heated to an internal temperature of 75 °C, while submissions replicating fish AORs were heated to an internal temperature of 63 °C.

Prior to advancing to sensory evaluation, all submissions underwent testing for common foodborne pathogens (*E. coli, L. monocytogenes, S. aureus,* and *Salmonella sp.*) at an accredited facility. Testing for these foodborne pathogens is a common requirement of national food regulations globally and was important to assert the safety of submissions post-production. One of the proposed advantages of meat alternatives is that, by decoupling meat production from animal agriculture, the risk of certain foodborne pathogens (e.g., *E. coli*) commonly found in animal gastrointestinal tracts should be significantly lower ^77^. These pathogens can also be introduced during manufacturing and handling so while the risk may be lower for meat alternatives compared to animal-based meat, testing for them will continue to be important.

The specific pathogen profile of meat alternatives has yet to be established so the FTNB framework included ACC testing to account for this. We anticipate refinement of the food safety testing portfolio as additional information emerges on food safety risks associated with meat alternatives.

### Safety concerns specific to cell-based meat production

There are limited examples of cell-based meat products being sold to the public and the regulatory framework is only now taking shape. There are also unique food safety concerns stemming from the expansion of animal cells *in vitro* that need to be considered. The FTNB framework incorporated a wide range of approaches for assessing food safety of cell-based products.

### Mycoplasma and other adventitious agents

Adventitious agents are broadly defined as bacteria, fungi, and viruses that can contaminate biomanufacturing processes. These can have a negative impact on yields and productivity but given the intended usage of cells grown for cell-based meat, adventitious agents represent a food safety concern. Proactive mitigation through good manufacturing processes and proper maintenance of sterility are at the forefront for reducing the risk of adventitious agents; however, formalizing and standardizing diagnostic testing will be an important control as adoption of cell-based foods increases.

Biocontrol procedures are well established in cell banking and were required of cells used in the competition ^78,79^. Cell banks were required to test for bacteria, viruses and *Mycoplasma* using real time PCR and through measurement of reverse transcriptase activity or other assays ^80^. In instances where commercial cell lines were utilized, competing teams were able to provide QC documentation from the supplier asserting acceptable results for parameters related to adventitious agents. Regular testing on working cell banks was required to ensure they remained free of *Mycoplasma* contamination by monitoring using PCR-based *Mycoplasma* detection kits and microscopy.

### Serum

Substantial efforts are underway to eliminate the reliance of cell-based meat production on animal sera ^81–83^; however, cell lines can be recalcitrant to serum-free adaptation and cell-based meat manufacturing can still incorporate serum. For FTNB, any serum used was required to be bovine spongiform encephalopathy/transmissible spongiform encephalopathy (BSE/TSE) certified. Transmissible encephalopathies are an enigmatic class of prion-based adventitious agents. Animal sera is a possible source of BSE/TSE, representing a food safety risk for cell-based foods ^84^. Requiring the use of TSE/BSE-Free serum was aimed at mitigating this risk as methods for detection are primarily designed to detect prions in tissues or tissue homogenates from livestock.

### Transparency and disclosure related to genetic modification

Genetic modification is a common strategy to establish immortalized cell lines ^85^ and has also been implemented as a tool to enhance the proliferation, nutritional properties ^86^, and amenability to serum-free growth ^87^ for cell lines with applications in cell-based meat manufacturing. There are a variety of approaches to genetic modification, many of which incorporate the use of exogenous genes that could create additional food safety concerns (e.g., SV-40-mediated immortalization ^88^). Genetic modification also introduces additional complexity from a regulatory standpoint, with certain countries prohibiting its use in food production.

XPRIZE FTNB strived to be solution agnostic and applied this policy to the use of genetic modification. Teams choosing to use genetically modified cell lines were required to disclose the nature of genetic modification to enable a risk-based assessment. Disclosure included: methodology used for genetic modification, specific genomic regions targeted, source(s) of any exogenous DNA, and supporting sequencing data which were reviewed by a panel of experts. No teams utilizing genetic modification proceeded to sensory evaluation.

### Cell line characterization – tumorigenicity and invasion studies using humanized mouse models

High rates of proliferation, anchorage-independent growth, and immortality are all highly desirable properties of cell lines intended for use in cell-based meat production. They are also hallmarks of oncogenic phenotypes ^89^, indicating rigorous cell line characterization is required before any cell lines are used for cell-based meat production. This is especially true of spontaneously immortalized cell lines, where the specific genetic changes that are responsible for immortalization can be difficult to pinpoint. A key outcome of the XPRIZE FTNB competition was the identification of a knowledge gap and the need for more research to enable data-driven risk assessment of meat alternatives produced using cell lines exhibiting these phenotypes. A FAO assessment found associated risks to be extremely low ^74^; however, until comprehensive data becomes available, empirical risk assessment is needed to proactively alleviate consumer concerns. As part of this risk assessment, the FTNB framework recommends characterizing cells for tumorigenicity and invasion using humanized mouse models ^90,91^. Tumorigenicity assays are routinely performed on master cell banks and should be part of the safety assessment for any cell lines used in the production of cell-based meat. Continued monitoring of master and working cell banks was recommended using adjunctive approaches such as high-throughput sequencing to characterize genetic stability.

### Cost Breakdown of the FTNB Framework

The FTNB framework was designed to be both modular and scalable, making it accessible to a wide range of potential end-users. Testing relies on well-established, standardized methodology that can be conducted in-house or by contracting a commercial testing lab. The cost of structural and physical characteristic testing and nutrition testing allows for the framework to be applied multiple times throughout product development, allowing for iterative improvement. Process level assessments and food safety are components of the framework that can be incorporated during later stages of product development, once a product and process has reached a level of maturity warranting such investment. A full cost breakdown of the FTNB framework is outlined in Table 3. These costs are reflective of testing conducted throughout FTNB for each competing team. Testing was conducted in the USA and UAE (see Methods for details) and actual costs may differ depending on location, service provider and scope of work.

**Table 3.** Cost breakdown of the FTNB framework (per product assessed)

| Testing Parameter | Cost (USD) |
| --- | --- |
| Structure and Physical Characteristics | \$1,500.00 |
| Nutrition | \$4,500.00 |
| Screening LCA | \$20,500.00 |
| TEA | \$8,000.00 |
| Sensory Evaluation | \$16,500.00 |
| Food Safety Testing | \$1,100.00 |
| **Total** | **\$52,100.00** |

## Discussion

Agricultural systems are increasingly strained which is exacerbating the sustainability challenges associated with traditional livestock farming, including deforestation, biodiversity loss, GHG emissions and antibiotic usage. The health and productivity of fisheries ^92,93^ and ocean communities are similarly challenged by increasing intensity of harvest and exploitation. A growing community of researchers have begun working on technology-based solutions to these challenges in the form of meat alternatives. There are a variety of approaches to producing meat alternatives, each with its own profile of proposed benefits and drawbacks. Studies focusing on consumer acceptance of meat alternatives suggest the market and potential for positive impact exist; however, they also indicate widespread consumer adoption is contingent on meat alternatives achieving near-parity with conventional meat in terms of taste, performance, and price ^94,95^.

XPRIZE Feed the Next Billion encouraged development of meat alternatives capable of replicating or outperforming conventional meat across a wide range of criteria. Evaluating the performance of competition submissions to determine if the standard of ‘replicate or outperform’ was achieved required the curation of a framework to systematically and holistically evaluate meat alternatives. We developed the framework (Table 4) to assess product and process dependent factors, allowing for comparison between meat alternatives and their AOR, as well as for direct comparisons between different alternative meat products. The framework is based on well established approaches to assessing meat quality and nutrition, allowing for testing results on meat alternatives to be meaningfully compared to the existing trove of meat science data available in academic literature. This feature of the framework was intentional and aimed at making the framework accessible to start ups, companies, researchers, investors, and any other stakeholders working in the field of alternative meat and cellular agriculture.

**Table 4.**
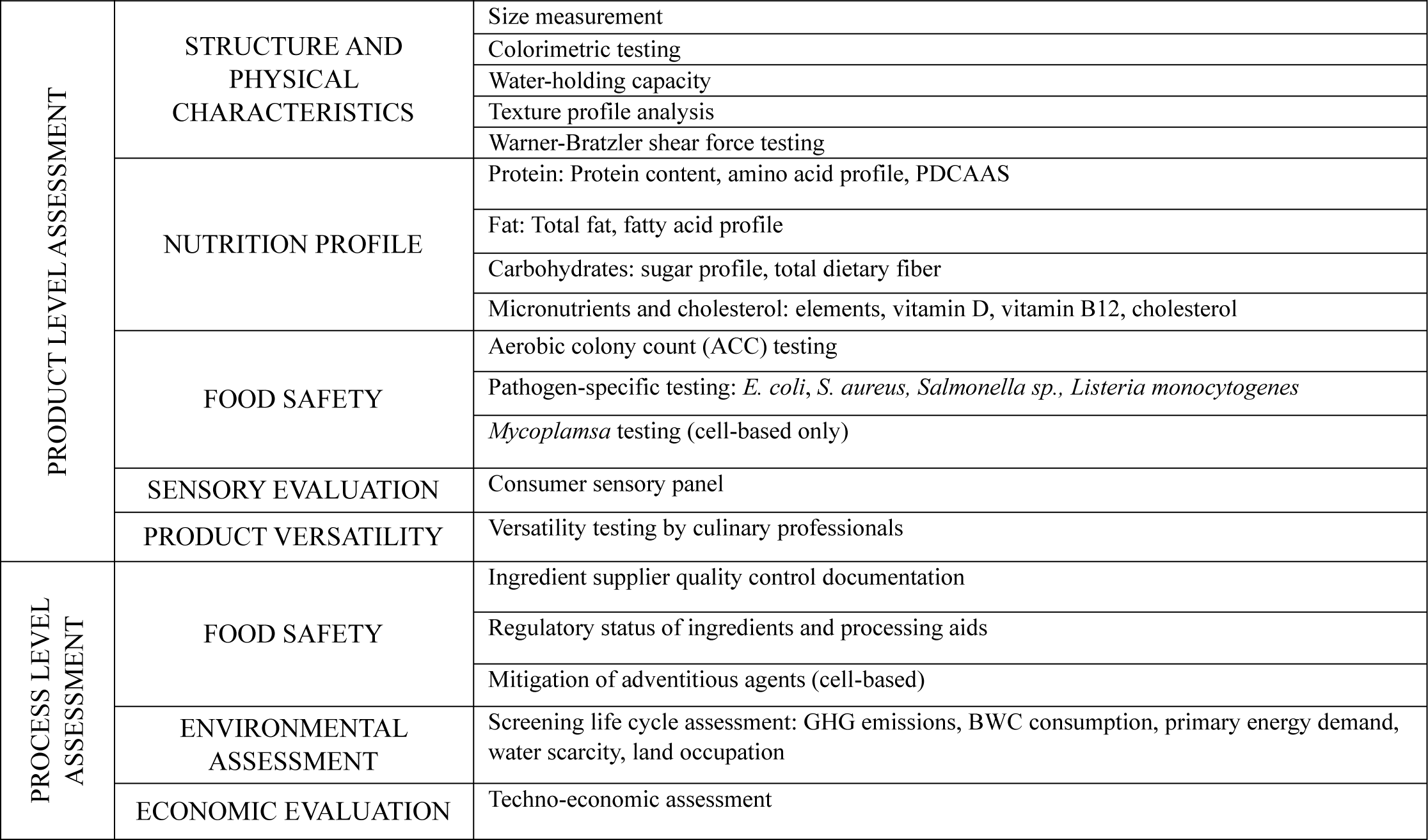
The XPRIZE Feed the Next Billion Framework.

The modular approach to constructing this framework and to assessing meat alternatives in comparison to their AOR allows for it to be incorporated as a part of iterative product development. Most of the testing focused on product level assessment could be incorporated during the initial stages of product development, allowing for short feedback loops to guide reformulation focused on optimizing for structure, nutrition, sensory and versatility. After achieving desired outcomes in these criteria, incorporating the process level aspects of the framework allow for key insight into product sustainability and unit economics.

The framework described here was used throughout the Feed the Next Billion competition, allowing for selection of teams advancing through various stages and ultimately to the competition finals. This test case suggested the framework enabled meaningful comparisons between meat alternatives and conventional meat. Correspondingly, we believe the framework represents a starting point and opportunity for the field of alternative proteins to move towards a more standardized approach to evaluation. We anticipate that the framework will be refined as additional data reveals which aspects have the strongest predictive power for consumer appeal and ultimately commercial success. Application of this framework and future versions of it will help improve data quality in the field, helping to better substantiate claims made regarding product quality, sustainability benefits, and economic viability.

## Methods

### Structure, physical characteristics and nutrition

Structural and physical characteristics and nutrition testing for all competition submissions and chicken and fish fillet references (termed ‘animal-origin references’ (AORs) was conducted by Eurofins National Food Lab USA. A list of methodology is included in Table 5.

**Table 5.**
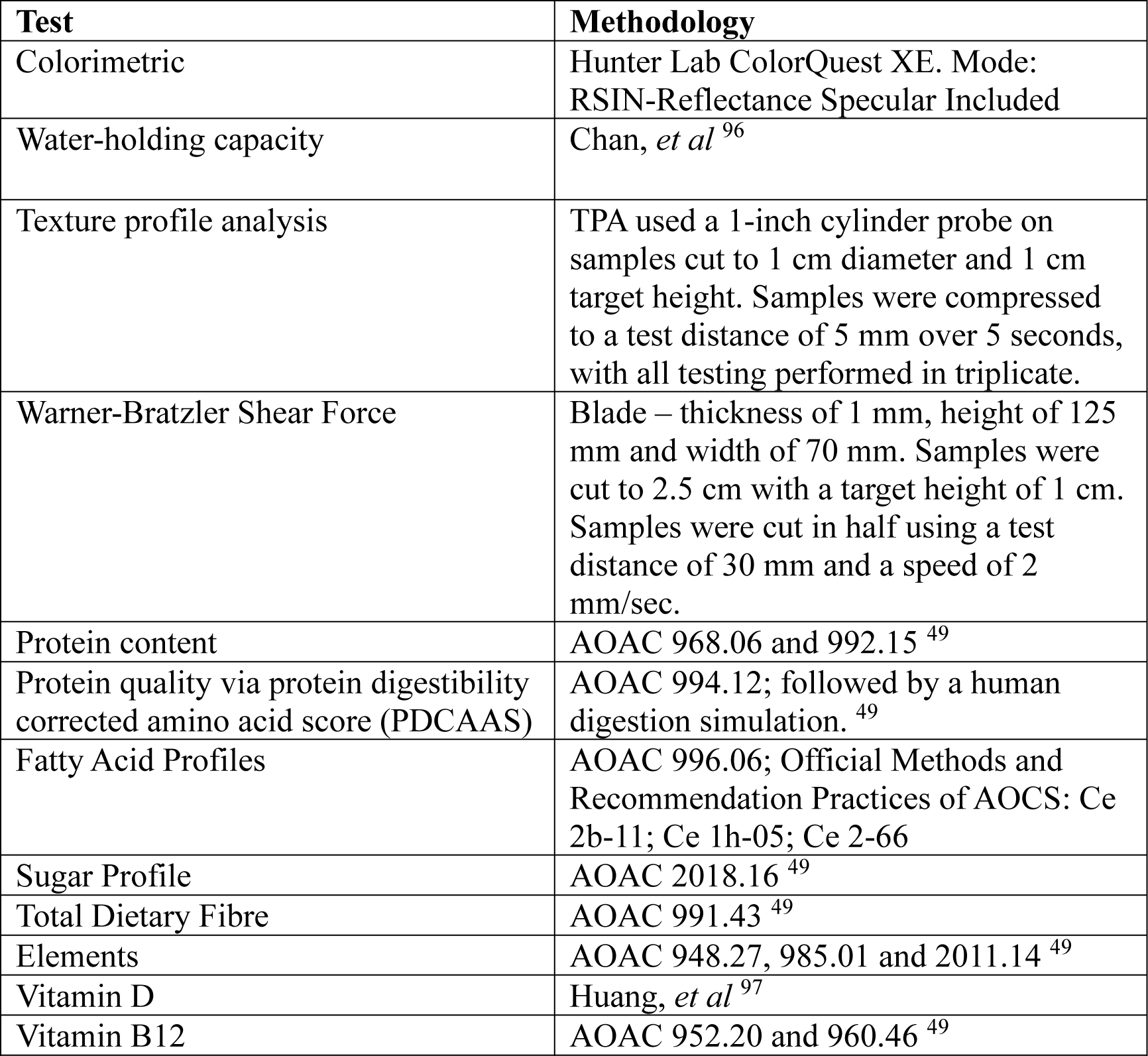
Testing methodology.

### Product versatility

Product versatility testing was performed by the International Center for Culinary Arts (ICCA) Abu Dhabi. Sixteen culinary professionals assessed submissions for: performance using a dry heat cooking method (e.g., grilling), performance using a moist heat cooking method (e.g., steaming), performance during dry heat cooking in combination with breading, texture after cooking with dry heat method, texture after cooking with moist heat method, texture after exposure to acid marination, and overall opinion of ease of use. Each assessment category was scored using a five-point scale (1 - poor; 5 - excellent).

### Sensory evaluation

Sensory evaluation was conducted using consumer sensory panels performed by Ipsos (UAE). Panels consisted of 100 consumers that were non-rejectors of AOR and willing to try meat alternatives ^98^. Blinded tasting proceeded via a sequential monadic approach ^99,100^. Consumers tasted samples (AOR and meat alternative) ‘as-is’, then prepared in a recipe. In between tastings, consumers were asked to cleanse their palate using unsalted crackers and water. After each tasting, consumers evaluated products for overall opinion, overall tase, overall aroma and overall texture using a seven-point scale (1 – very poor; 7 – excellent). Purchase intent data was also collected using a five-point scale (1 – not likely to purchase; 5 – very likely to purchase). Purchase intent data was for research purposes only and not used for evaluating competitors in FTNB.

### Environmental assessment

To assess the environmental impacts from production of competition submissions, screening level life cycle assessments (LCAs) were performed by WSP USA Inc. in alignment with ISO standards 14040 and 14044. The goals of the screening LCA were to assess the GHG emissions, land occupation, water use, water scarcity and energy use from cradle-to-gate of 1 kg of product. GHG emissions were assessed including their contribution to global warming potential over a 100-year period, reported as kgCO2e. Energy use was assessed as primary energy demand (PED) from non-renewable resources and expressed in megajoules. Blue water consumption (BWC) measured the amount of surface or ground water consumed for production, while water scarcity was assessed using the AWARE method ^101^.

### Economic evaluation

Economic feasibility of FTNB entrants was assessed using a techno-economic assessment and business case analysis developed and performed Pi Innovations (USA). A list of parameters analyzed is included in Table 6.

**Table 6.**
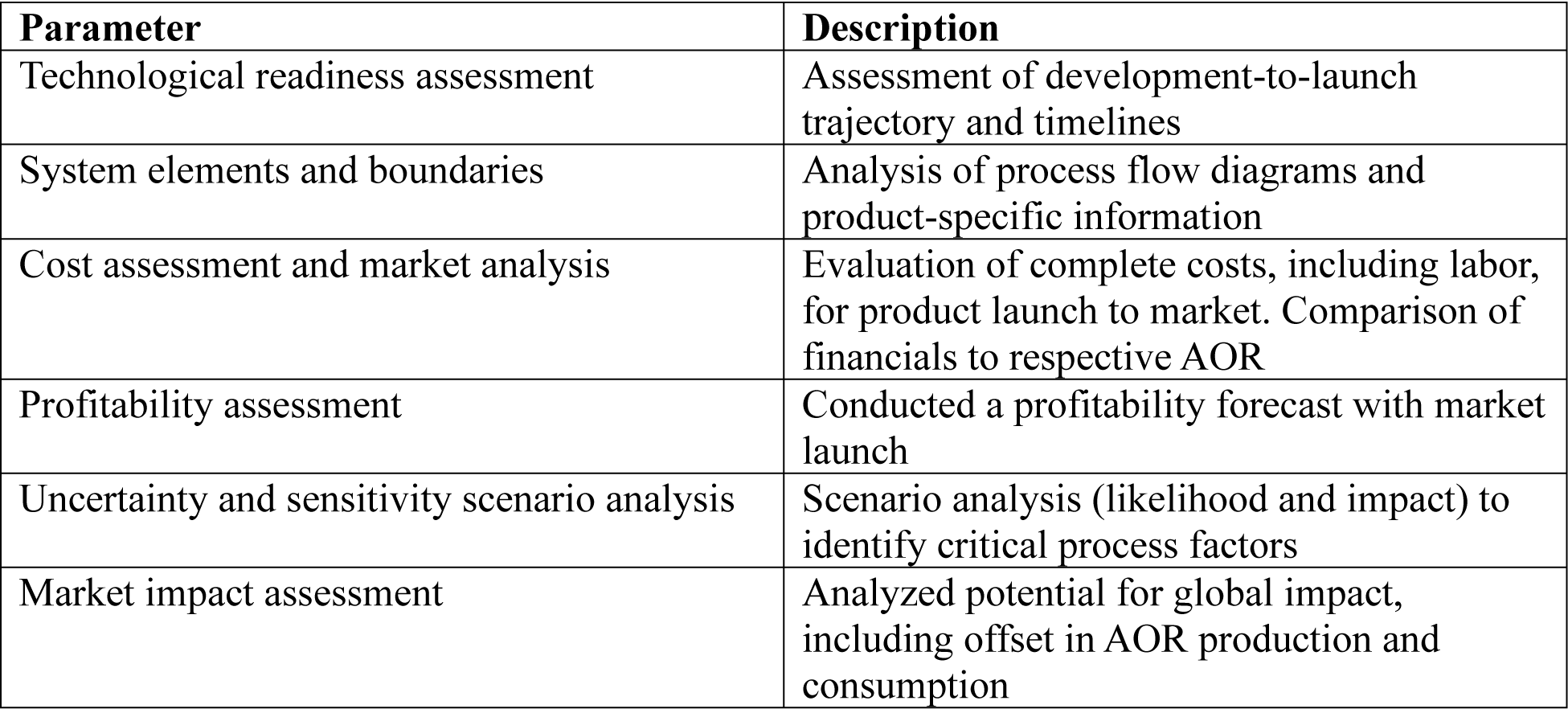
Economic and business case parameters evaluated.

### Food Safety

Food safety testing was performed by the Abu Dhabi Quality & Conformity Council Testing Laboratory. Aerobic colony count, *Escherichia coli* and *Staphylococcus aureus* testing were performed by using Petrifilm ^102–104^. Testing for *Listeria monocytogenes* and *Salmonella sp.* was via Vitek immunodiagnostic assay (VIDAS) ^105,106^.

## Acknowledgements

We thank all teams and competitors that participated in XPRIZE Feed the Next Billion. Our gratitude extends to the Competition Judging Panel, including Brian Jacobson and Keith Cox, Advisory Board, as well as the testing partners and vendors that implemented the various assessment components of the framework. We acknowledge the generous support of the Competition sponsors: ASPIRE, TRF and an anonymous donor. Figure 1 was made using BioRender.

## References

1. Ritchie H, Rosado P, Roser M. Meat and dairy production. Our world in data. 2024.

2. Garcia SN, Osburn BI, Jay-Russell MT. One Health for Food Safety, Food Security, and Sustainable Food Production. Frontiers in Sustainable Food Systems. 2020;4.

3. Zhang T, Nickerson R, Zhang W, et al. The impacts of animal agriculture on One Health—Bacterial zoonosis, antimicrobial resistance, and beyond. One Health. 2024;18:100748.

4. Eisfeld AJ, Biswas A, Guan L, et al. Pathogenicity and transmissibility of bovine H5N1 influenza virus. Nature. 2024;633(8029):426-432.

5. MacLeod MJ, Vellinga T, Opio C, et al. Invited review: A position on the Global Livestock Environmental Assessment Model (GLEAM). Animal. 2018;12(2):383–397.

6. Xu X, Sharma P, Shu S, et al. Global greenhouse gas emissions from animal-based foods are twice those of plant-based foods. Nature Food. 2021;2(9):724–732.

7. Eisen MB, Brown PO. Rapid global phaseout of animal agriculture has the potential to stabilize greenhouse gas levels for 30 years and offset 68 percent of CO2 emissions this century. PLOS Climate. 2022;1(2):e0000010.

8. UN. Paris Agreement. Article 2*(a).* 2015.

9. Sylvester JM, Gutiérrez-Zapata DM, Pérez-Marulanda L, et al. Analysis of food system drivers of deforestation highlights foreign direct investments and urbanization as threats to tropical forests. Scientific Reports. 2024;14(1):15179.

10. Gerbens-Leenes PW, Mekonnen MM, Hoekstra AY. The water footprint of poultry, pork and beef: A comparative study in different countries and production systems. Water Resources and Industry. 2013;1-2:25–36.

11. Dudley N, Alexander S. Agriculture and biodiversity: a review. Biodiversity. 2017;18(2-3):45–49.

12. Tuomisto HL, Teixeira de Mattos MJ. Environmental Impacts of Cultured Meat Production. Environmental Science & Technology. 2011;45(14):6117–6123.

13. Smetana S, Ristic D, Pleissner D, Tuomisto HL, Parniakov O, Heinz V. Meat substitutes: Resource demands and environmental footprints. Resour Conserv Recycl. 2023;190:106831.

14. Rubio NR, Xiang N, Kaplan DL. Plant-based and cell-based approaches to meat production. Nat Commun. 2020;11(1):6276.

15. He J, Evans NM, Liu H, Shao S. A review of research on plant-based meat alternatives: Driving forces, history, manufacturing, and consumer attitudes. Comprehensive Reviews in Food Science and Food Safety. 2020;19(5):2639–2656.

16. Laestadius LI, Caldwell MA. Is the future of meat palatable? Perceptions of in vitro meat as evidenced by online news comments. Public Health Nutr. 2015;18(13):2457–2467.

17. Stevens H, Ruperti Y. Smart food: novel foods, food security, and the Smart Nation in Singapore. *Food*, Culture & Society. 2024;27(3):754–774.

18. Humbird D. Scale-up economics for cultured meat. Biotechnol Bioeng. 2021;118(8):3239–3250.

19. Negulescu PG, Risner D, Spang ES, et al. Techno-economic modeling and assessment of cultivated meat: Impact of production bioreactor scale. Biotechnology and Bioengineering. 2023;120(4):1055–1067.

20. Kerslake E, Kemper JA, Conroy D. What’s your beef with meat substitutes? Exploring barriers and facilitators for meat substitutes in omnivores, vegetarians, and vegans. Appetite. 2022;170:105864.

21. Hall C. Even after $1.6B in VC money, the lab-grown meat industry is facing ’massive’ issues 2024. https://techcrunch.com/2024/08/04/even-after-1-6b-in-vc-money-the-lab-grown-meat-industry-is-facing-massive-issues/.

22. Brehaut L. Plant-based meat sales fall ’significantly’ for second year in a row 2024. https://nationalpost.com/life/food/plant-based-vegan-meat-sales-fall-significantly-for-second-year-in-a-row.

23. Lopez-Pedrouso M, Rodriguez-Vazquez R, Purrinos L, et al. Sensory and Physicochemical Analysis of Meat from Bovine Breeds in Different Livestock Production Systems, Pre-Slaughter Handling Conditions and Ageing Time. Foods. 2020;9(2).

24. Dekkers BL, Boom RM, van der Goot AJ. Structuring processes for meat analogues. Trends in Food Science & Technology. 2018;81:25–36.

25. Varayil H, Meena, D., Mitra, J. . Plant-Based Foods: Advanced Structuring Techniques In: Structured Foods Taylor & Francis Group 2024:19.

26. Schätzlein E, Blaeser A. Recent trends in bioartificial muscle engineering and their applications in cultured meat, biorobotic systems and biohybrid implants. Communications Biology. 2022;5(1):737.

27. Scozzafava G, Corsi AM, Casini L, Contini C, Loose SM. Using the animal to the last bit: Consumer preferences for different beef cuts. Appetite. 2016;96:70–79.

28. Tanaka RI, Sakaguchi K, Yoshida A, Takahashi H, Haraguchi Y, Shimizu T. Production of scaffold-free cell-based meat using cell sheet technology. NPJ Sci Food. 2022;6(1):41.

29. Kawecki NS, Norris SCP, Xu Y, et al. Engineering multicomponent tissue by spontaneous adhesion of myogenic and adipogenic microtissues cultured with customized scaffolds. Food Res Int. 2023;172:113080.

30. Savadkoohi S, Hoogenkamp H, Shamsi K, Farahnaky A. Color, sensory and textural attributes of beef frankfurter, beef ham and meat-free sausage containing tomato pomace. Meat Sci. 2014;97(4):410–418.

31. Alfnes F, Atle GG, Gro S, Kari K. Consumers’ Willingness to Pay for the Color of Salmon: A Choice Experiment with Real Economic Incentives. American Journal of Agricultural Economics. 2006;88(4):1050–1061.

32. Ruedt C, Gibis M, Weiss J. Meat color and iridescence: Origin, analysis, and approaches to modulation. Comprehensive Reviews in Food Science and Food Safety. 2023;22(4):3366–3394.

33. Storebakken T, Foss P, Schiedt K, Austreng E, Liaaen-Jensen S, Manz U. Carotenoids in diets for salmonids: IV. Pigmentation of Atlantic salmon with astaxanthin, astaxanthin dipalmitate and canthaxanthin. Aquaculture. 1987;65(3):279–292.

34. Buttle L, Crampton V, Williams P. The effect of feed pigment type on flesh pigment deposition and colour in farmed Atlantic salmon, Salmo salar L. Aquaculture Research. 2001;32(2):103–111.

35. Suman SP, Nair MN, Joseph P, Hunt MC. Factors influencing internal color of cooked meats. Meat Science. 2016;120:133–144.

36. Warner RD. Chapter 14 - The eating quality of meat: IV—Water holding capacity and juiciness. In: Toldrá F, ed. Lawrie’s Meat Science (Ninth Edition). Woodhead Publishing; 2023:457–508.

37. Cheng Q, Sun DW. Factors affecting the water holding capacity of red meat products: a review of recent research advances. Crit Rev Food Sci Nutr. 2008;48(2):137–159.

38. Nishinari K, Kohyama K, Kumagai H, Funami T, Bourne MC. Parameters of Texture Profile Analysis. Food Science and Technology Research. 2013;19(3):519–521.

39. de Huidobro FR, Miguel E, Blázquez B, Onega E. A comparison between two methods (Warner–Bratzler and texture profile analysis) for testing either raw meat or cooked meat. Meat Science. 2005;69(3):527–536.

40. Caine WR, Aalhus JL, Best DR, Dugan MER, Jeremiah LE. Relationship of texture profile analysis and Warner-Bratzler shear force with sensory characteristics of beef rib steaks. Meat Science. 2003;64(4):333–339.

41. Herrero AM, de la Hoz L, Ordóñez JA, Herranz B, Romero de Ávila MD, Cambero MI. Tensile properties of cooked meat sausages and their correlation with texture profile analysis (TPA) parameters and physico-chemical characteristics. Meat Science. 2008;80(3):690–696.

42. Souppez J-BRG, Dages BAS, Pavar GS, Fabian J, Thomas JM, Theodosiou E. Mechanical properties and texture profile analysis of beef burgers and plant-based analogues. Journal of Food Engineering. 2025;385:112259.

43. Paredes J, Cortizo-Lacalle D, Imaz AM, Aldazabal J, Vila M. Application of texture analysis methods for the characterization of cultured meat. Sci Rep. 2022;12(1):3898.

44. Purohit AS, Reed C, Mohan A. Development and evaluation of quail breakfast sausage. LWT - Food Science and Technology. 2016;69:447–453.

45. Nasrollahzadeh F, Alexi N, Skov KB, Roman L, Sfyra K, Martinez MM. Texture profiling of muscle meat benchmarks and plant-based analogues: An instrumental and sensory design approach with focus on correlations. Food Hydrocolloids. 2024;151:109829.

46. Novaković S, Tomašević I. A comparison between Warner-Bratzler shear force measurement and texture profile analysis of meat and meat products: a review. IOP Conference Series: Earth and Environmental Science. 2017;85(1):012063.

47. Zupo V, Graber G, Kamel S, et al. Mercury accumulation in freshwater and marine fish from the wild and from aquaculture ponds. Environmental Pollution. 2019;255:112975.

48. Li J, Silver C, Gómez MI, Milstein M, Sogari G. Factors influencing consumer purchase intent for meat and meat substitutes. Future Foods. 2023;7:100236.

49. Horwitz W, Latimer GW. Official methods of analysis of AOAC International. 18th ed. Gaithersburg, Md.: AOAC International; 2005.

50. Schaafsma G. The protein digestibility-corrected amino acid score. J Nutr. 2000;130(7):1865S–1867S.

51. Boye J, Wijesinha-Bettoni R, Burlingame B. Protein quality evaluation twenty years after the introduction of the protein digestibility corrected amino acid score method. British Journal of Nutrition. 2012;108(S2):S183–S211.

52. Hertzler SR, Lieblein-Boff JC, Weiler M, Allgeier C. Plant Proteins: Assessing Their Nutritional Quality and Effects on Health and Physical Function. Nutrients. 2020;12(12).

53. Dimina L, Rémond D, Huneau J-F, Mariotti F. Combining Plant Proteins to Achieve Amino Acid Profiles Adapted to Various Nutritional Objectives—An Exploratory Analysis Using Linear Programming. Frontiers in Nutrition. 2022;8.

54. Ruxton CHS. The benefits of fish consumption. Nutrition Bulletin. 2011;36(1):6–19.

55. Deng Q, Wang Z, Fu L, et al. High-moisture extrusion of soy protein: Effects of insoluble dietary fiber on anisotropic extrudates. Food Hydrocolloids. 2023;141:108688.

56. Samtiya M, Aluko RE, Dhewa T, Moreno-Rojas JM. Potential Health Benefits of Plant Food-Derived Bioactive Components: An Overview. Foods. 2021;10(4).

57. Messina M, Duncan AM, Glenn AJ, Mariotti F. Perspective: Plant-Based Meat Alternatives Can Help Facilitate and Maintain a Lower Animal to Plant Protein Intake Ratio. Adv Nutr. 2023;14(3):392–405.

58. Estell M, Hughes J, Grafenauer S. Plant Protein and Plant-Based Meat Alternatives: Consumer and Nutrition Professional Attitudes and Perceptions. Sustainability. 2021;13(3).

59. Romão B, Botelho RB, Torres ML, et al. Nutritional Profile of Commercialized Plant-Based Meat: An Integrative Review with a Systematic Approach. Foods. 2023;12(3).

60. Alessandrini R, Brown MK, Pombo-Rodrigues S, Bhageerutty S, He FJ, MacGregor GA. Nutritional Quality of Plant-Based Meat Products Available in the UK: A Cross-Sectional Survey. Nutrients. 2021;13(12).

61. Zimmermann MB, Hurrell RF. Nutritional iron deficiency. The Lancet. 2007;370(9586):511–520.

62. Shlisky J, Mandlik R, Askari S, et al. Calcium deficiency worldwide: prevalence of inadequate intakes and associated health outcomes. Annals of the New York Academy of Sciences. 2022;1512(1):10–28.

63. Allen LH. Vitamin B-12. Adv Nutr. 2012;3(1):54–55.

64. Ozturk OK, Hamaker BR. Texturization of plant protein-based meat alternatives: Processing, base proteins, and other constructional ingredients. Future Foods. 2023;8:100248.

65. Russo Spena S, Pasquino R, Sarrica A, Delmonte M, Yang C, Grizzuti N. Kinetics of acid hydrolysis of k-Carrageenan by in situ rheological follow-up. Food Hydrocolloids. 2023;144:108953.

66. Giacalone D, Clausen MP, Jaeger SR. Understanding barriers to consumption of plant-based foods and beverages: insights from sensory and consumer science. Current Opinion in Food Science. 2022;48:100919.

67. Shanmugam K, Bryngelsson S, Östergren K, Hallström E. Climate Impact of Plant-based Meat Analogues: A Review of Life Cycle Assessments. Sustainable Production and Consumption. 2023;36:328–337.

68. Trinidad KR, Ashizawa R, Nikkhah A, et al. Environmental life cycle assessment of recombinant growth factor production for cultivated meat applications. Journal of Cleaner Production. 2023;419:138153.

69. de Vries M, de Boer IJM. Comparing environmental impacts for livestock products: A review of life cycle assessments. Livestock Science. 2010;128(1):1–11.

70. Rust JM. The impact of climate change on extensive and intensive livestock production systems. Animal Frontiers. 2019;9(1):20–25.

71. Smetana S, Ristic D, Pleissner D, Tuomisto HL, Parniakov O, Heinz V. Meat substitutes: Resource demands and environmental footprints. Resources, Conservation and Recycling. 2023;190:106831.

72. Tuomisto HL, Allan SJ, Ellis MJ. Prospective life cycle assessment of a bioprocess design for cultured meat production in hollow fiber bioreactors. Science of The Total Environment. 2022;851:158051.

73. Sinke P, Swartz E, Sanctorum H, van der Giesen C, Odegard I. Ex-ante life cycle assessment of commercial-scale cultivated meat production in 2030. The International Journal of Life Cycle Assessment. 2023;28(3):234–254.

74. FAO & WHO. Food safety aspects of cell-based food. 2023.

75. Goodman RE, Ebisawa M, Ferreira F, et al. AllergenOnline: A peer-reviewed, curated allergen database to assess novel food proteins for potential cross-reactivity. Molecular Nutrition & Food Research. 2016;60(5):1183–1198.

76. Lin X, Duan N, Wu J, Lv Z, Wang Z, Wu S. Potential food safety risk factors in plant-based foods: Source, occurrence, and detection methods. Trends in Food Science & Technology. 2023;138:511–522.

77. Ong KJ, Johnston J, Datar I, Sewalt V, Holmes D, Shatkin JA. Food safety considerations and research priorities for the cultured meat and seafood industry. Comprehensive Reviews in Food Science and Food Safety. 2021;20(6):5421–5448.

78. Andrews PW, Baker D, Benvinisty N, et al. Points to Consider in the Development of Seed Stocks of Pluripotent Stem Cells for Clinical Applications: International Stem Cell Banking Initiative (ISCBI). Regenerative Medicine. 2015;10(sup2):1–44.

79. Stacey GN, Masters JR. Cryopreservation and banking of mammalian cell lines. Nature Protocols. 2008;3(12):1981–1989.

80. Barone PW, Wiebe ME, Leung JC, et al. Viral contamination in biologic manufacture and implications for emerging therapies. Nature Biotechnology. 2020;38(5):563–572.

81. Skrivergaard S, Young JF, Sahebekhtiari N, et al. A simple and robust serum-free media for the proliferation of muscle cells. Food Res Int. 2023;172:113194.

82. Venkatesan M, Semper C, Skrivergaard S, et al. Recombinant production of growth factors for application in cell culture. iScience. 2022;25(10):105054.

83. Stout AJ, Mirliani AB, Rittenberg ML, et al. Simple and effective serum-free medium for sustained expansion of bovine satellite cells for cell cultured meat. Commun Biol. 2022;5(1):466.

84. Chou ML, Bailey A, Avory T, Tanimoto J, Burnouf T. Removal of transmissible spongiform encephalopathy prion from large volumes of cell culture media supplemented with fetal bovine serum by using hollow fiber anion-exchange membrane chromatography. PLoS One. 2015;10(4):e0122300.

85. Stout AJ, Arnett MJ, Chai K, et al. Immortalized Bovine Satellite Cells for Cultured Meat Applications. ACS Synthetic Biology. 2023;12(5):1567–1573.

86. Stout AJ, Mirliani AB, Soule-Albridge EL, Cohen JM, Kaplan DL. Engineering carotenoid production in mammalian cells for nutritionally enhanced cell-cultured foods. Metabolic Engineering. 2020;62:126–137.

87. Stout AJ, Zhang X, Letcher SM, et al. Engineered autocrine signaling eliminates muscle cell FGF2 requirements for cultured meat production. bioRxiv. 2023.

88. Jha KK, Banga S, Palejwala V, Ozer HL. SV40-Mediated immortalization. Exp Cell Res. 1998;245(1):1–7.

89. Mori S, Chang JT, Andrechek ER, et al. Anchorage-independent cell growth signature identifies tumors with metastatic potential. Oncogene. 2009;28(31):2796–2805.

90. Rampetsreiter P, Casanova E, Eferl R. Genetically modified mouse models of cancer invasion and metastasis. Drug Discovery Today: Disease Models. 2011;8(2):67–74.

91. Yin L, Wang XJ, Chen DX, Liu XN, Wang XJ. Humanized mouse model: a review on preclinical applications for cancer immunotherapy. Am J Cancer Res. 2020;10(12):4568–4584.

92. Costello C, Cao L, Gelcich S, et al. The future of food from the sea. Nature. 2020;588(7836):95–100.

93. Sumaila UR, Tai TC. End overfishing and increase the resilience of the ocean to climate change. Frontiers in Marine Science. 2020;7:523.

94. Onwezen MC, Bouwman EP, Reinders MJ, Dagevos H. A systematic review on consumer acceptance of alternative proteins: Pulses, algae, insects, plant-based meat alternatives, and cultured meat. Appetite. 2021;159:105058.

95. Siddiqui SA, Alvi T, Sameen A, et al. Consumer Acceptance of Alternative Proteins: A Systematic Review of Current Alternative Protein Sources and Interventions Adapted to Increase Their Acceptability. Sustainability. 2022;14(22).

96. Chan SS, Roth B, Jessen F, Jakobsen AN, Lerfall J. Water holding properties of Atlantic salmon. Compr Rev Food Sci Food Saf. 2022;21(1):477–498.

97. Huang M, LaLuzerne P, Winters D, Sullivan D. Measurement of vitamin D in foods and nutritional supplements by liquid chromatography/tandem mass spectrometry. J AOAC Int. 2009;92(5):1327–1335.

98. Rogers L. Sensory panel management: a practical handbook for recruitment, training and performance. Woodhead Publishing; 2017.

99. Colyar JM, Eggett DL, Steele FM, Dunn ML, Ogden LV. Sensitivity comparison of sequential monadic and side-by-side presentation protocols in affective consumer testing. Journal of food science. 2009;74(7):S322–S327.

100. Komanska H. Sequential monadic designs: some theoretical considerations. Journal of sensory studies. 1990;4(3):201–211.

101. Boulay A-M, Bare J, Benini L, et al. The WULCA consensus characterization model for water scarcity footprints: assessing impacts of water consumption based on available water remaining (AWARE). The International Journal of Life Cycle Assessment. 2018;23(2):368–378.

102. Knight MT, Newman MC, Benzinger MJ, Jr., et al. Comparison of the Petrifilm dry rehydratable film and conventional culture methods for enumeration of yeasts and molds in foods: collaborative study. J AOAC Int. 1997;80(4):806–823.

103. Bird P, Flannery J, Crowley E, Agin J, Goins D, Jechorek R. Evaluation of the 3M Petrifilm Rapid Yeast and Mold Count Plate for the Enumeration of Yeast and Mold in Food: Collaborative Study, First Action 2014.05. J AOAC Int. 2015;98(3):767–783.

104. Kinneberg KM, Lindberg KG. Dry rehydratable film method for rapid enumeration of coliforms in foods (3M Petrifilm Rapid Coliform Count plate): collaborative study. J AOAC Int. 2002;85(1):56–71.

105. Curiale MS, Gangar V, Gravens C. VIDAS enzyme-linked fluorescent immunoassay for detection of Salmonella in foods: collaborative study. J AOAC Int. 1997;80(3):491–504.

106. Crowley E, Bird P, Flannery J, et al. Evaluation of VIDAS UP Listeria assay (LPT) for the detection of Listeria in a variety of foods and environmental surfaces: First Action 2013.10. J AOAC Int. 2014;97(2):431–441.

